# An Intrinsically Disordered Peptide Tag that Confers an Unusual Solubility to Aggregation-Prone Proteins

**DOI:** 10.1101/2021.08.05.455358

**Authors:** Byung Hoon Jo

**Author notes:** Correspondence to B. H. Jo.

## Abstract

There is a high demand for the production of recombinant proteins in *Escherichia coli* for biotechnological applications but their production is still limited by their insolubility. Fusion tags have been successfully used to enhance the solubility of aggregation-prone proteins; however, smaller and more powerful tags are desired for increasing the yield and quality of target proteins. Herein, NEXT tag, a 53 amino acid-length solubility enhancer, is described. The NEXT tag showed outstanding ability to improve both *in vivo* and *in vitro* solubilities with minimal effect on passenger proteins. The C-terminal region of the tag was mostly responsible for *in vitro* solubility, while the N-terminal region was essential for *in vivo* soluble expression. The NEXT tag appeared to be intrinsically disordered and seemed to exclude neighboring molecules and prevent protein aggregation by acting as an entropic bristle. This novel peptide tag should have general use as a fusion partner to increase the yield and quality of difficult-to-express proteins.

**IMPORTANCE:** Production of recombinant protein in *Escherichia coli* still suffers from the insolubility problem. Conventional solubility enhancers with large sizes represented by maltose-binding protein (MBP) have remained as the first-choice tags, however, the success in the soluble expression of tagged protein is largely unpredictable. In addition, the large tags can negatively affect the function of target proteins. In this work, NEXT tag, an intrinsically disordered peptide, was introduced as a small but powerful alternative to MBP. The NEXT tag could significantly improve both expression level and solubility of target proteins including a thermostable carbonic anhydrase and a polyethylene terephthalate (PET)-degrading enzyme that are remarkable enzymes for environmental bioremediation.

## INTRODUCTION

Recombinant protein expression underpins protein engineering and metabolic pathway engineering for the production of biotherapeutics, bioreporters, biocatalysts, and various industrial chemicals as well as biochemical studies of protein. Bacterial hosts such as *Escherichia coli* have remained the preferred hosts for recombinant protein expression due to their fast growth rate to high cell densities, ease of genetic manipulation, and scale-up simplicity (1). However, the high rate of protein synthesis/folding and high-level accumulation of heterologous protein in *E. coli* often lead to the formation of inclusion bodies that are intracellular aggregates of misfolded, partially folded, or even fully folded proteins, limiting the yield of recombinant protein (1-3).

Fusion protein tags have been widely used as effective solubility enhancers for aggregation-prone proteins. When fused to the N termini of target proteins, these solubility tags not only improve the solubility of passenger proteins but also increase their expression level by providing sequence contexts for more efficient translation initiation (4). Despite numerous examples of their applications, the successful selection of effective tags for a given target protein still relies heavily on a trial-and-error approach. Although maltose-binding protein (MBP) and N-utilization substance A (NusA) are the best working tags (4, 5), they are relatively large proteins (> 40 kDa) that can impart a higher metabolic burden than smaller tags and increase the chance of full-length proteins undergoing incomplete synthesis (6), potentially leading to a lower yield of target protein. In addition, while fusion tags are generally removed by chemical or enzymatic cleavage after soluble expression and purification, proteins that have intrinsically poor solubilities need to be used in their tagged forms since they precipitate after the solubility tags are removed (1, 5, 7). In this case, smaller tags are more desirable to minimize the effect of fusion tags on the inherent properties of passenger proteins (7, 8). Collectively, there is still considerable demand to expand the repertoire of solubility tags that are more powerful but smaller than conventional tags.

The carbonic anhydrase of the marine bacterium *Hydrogenovibrio marinus* (*hm*CA) is a highly soluble protein (9). *hm*CA contains an unusual N-terminal extension that is not essential for its catalytic function. When the N-terminal sequence was truncated, the solubility of recombinant *hm*CA was drastically reduced, and it was expressed mostly in an insoluble form. Inspired by this observation, it was hypothesized that the 53 amino acid-length N-terminal extension sequence (designated as NEXT) could be used as a fusion tag to improve solubility. In this study, it was demonstrated that the small NEXT tag is an intrinsically disordered protein that is exceptionally powerful for improving the solubility and expression level of passenger proteins with minimal influence on the target protein.

## RESULTS AND DISCUSSION

### Effect of the fusion tags on the *in vivo* solubility of recombinant proteins

By convention, there are two types of protein solubilities: *in vivo* solubility and *in vitro* solubility (10). Low *in vivo* protein solubility is often observed when a recombinant protein is overexpressed in a bacterial host, generally resulting in the formation of inclusion bodies (1, 2). When a protein has low *in vitro* solubility, aggregates can be formed even with a folded, isolated protein (10, 11). In this case, the protein can remain soluble only at a low concentration.

To evaluate the efficiency of the NEXT tag in improving *in vivo* solubility, the tag was fused to the N termini of several difficult-to-express proteins via a flexible linker **(Fig. 1a)**. The passenger proteins were selected based on their potential applications: human epidermal growth factor (hEGF, 6.4 kDa) as a therapeutic protein, green fluorescent protein (GFP, 26.9 kDa) as a bioreporter, and carbonic anhydrase from *Thermovibrio ammonificans* (*ta*CA, 25.9 kDa) and polyethylene terephthalate (PET)-hydrolyzing enzyme from *Ideonella sakaiensis* (*is*PETase, 27.7 kDa) as biocatalysts for bioremediation. Other commonly used solubility tags, including MBP (12), glutathione S-transferase (GST) (13), and an 8-kDa protein from *Fasciola hepatica* (Fh8) (4), were also tested for comparison **(Table 1)**. The fusion proteins were constructed, and their expression patterns were analyzed after soluble/insoluble fractionation.

**TABLE 1.** Properties of solubility tags used in this study

|  | Amino acid | Molecular | Phosphorylation | pI | Net charge | Mean |
| --- | --- | --- | --- | --- | --- | --- |
|  | length | weight (kDa) | sites |  |  | hydropathy |
| MBP | 367 | 40.4 | 26 | 5.1 | -10 | 0.465 |
| GST | 220 | 25.7 | 12 | 5.9 | -4 | 0.460 |
| Fh8 | 69 | 7.7 | 3 | 5.6 | -2 | 0.426 |
| NEXT | 53 | 5.5 | 0 | 8.1 | +1 | 0.397 |

**FIG 1.**
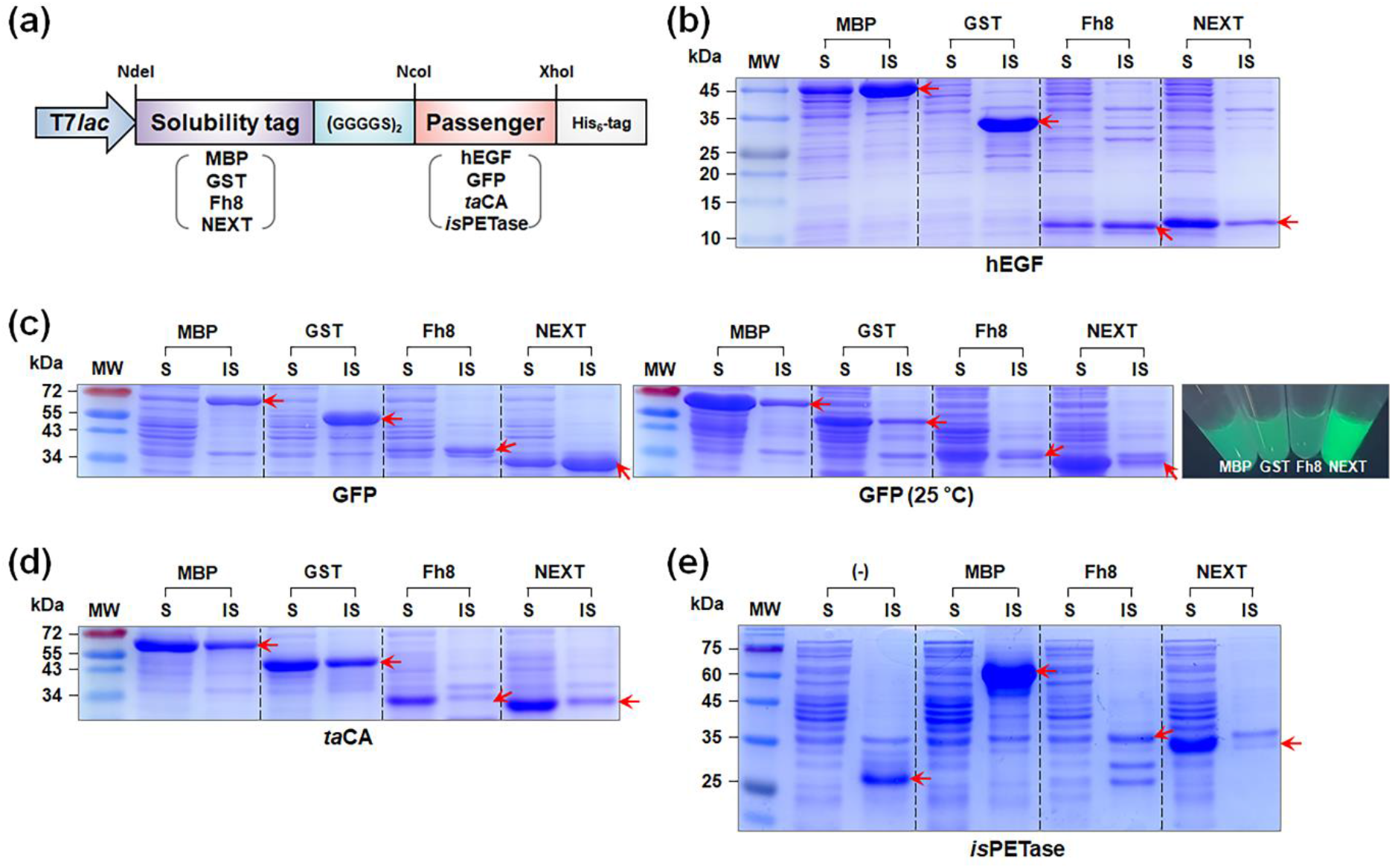
*In vivo* solubility of recombinant proteins fused with different solubility tags. (a) Expression cassettes of fusion proteins. Protein expression is under the control of the IPTG-inducible T7*lac* promoter. The passenger proteins are fused with the N-terminal solubility tags via the flexible (GGGGS)_2_ linker and with the C-terminal His_6_-tag. Coomassie blue-stained SDS-PAGE gel for the analysis of the expression of (b) hEGF, (c) GFP, (d) *ta*CA, and (e) *is*PETase fused with the different solubility tags at 37 °C or 25 °C (only for GFP). The fluorescence image in (c) is for GFP at 25 °C. The arrow indicates the band position of each recombinant protein. Lane: MW, molecular mass marker; S, soluble fraction; IS, insoluble fraction.

hEGF is a protein hormone that can be used as a wound healing agent by stimulating epidermal regeneration (14). Although the production of hEGF in bacterial cells has been reported, high-level soluble expression of hEGF in *E. coli* is still challenging. The expression level of soluble NEXT-hEGF was the highest among the tested constructs, and the percentage of the soluble fraction was 87 ± 9%, even at 37 °C **(Fig. 1b)**, which was also one of the highest values ever reported (15-17). GFP is the most popular fluorescent protein for bioimaging and sensing applications (18). When GFP was fused to the solubility tags, a notable amount of soluble expression was not observed at 37 °C with all of the tested fusion proteins except for NEXT-GFP **(Fig. 1c)**. At 25 °C, however, all of the constructs showed high *in vivo* solubility, and again, the most remarkable soluble expression was that of NEXT-GFP, as its fluorescence was the brightest **(Fig. 1c)**. *ta*CA is one of the most thermostable carbonic anhydrases and has potential applications in bioinspired CO_2_ capture and utilization (19-21). Despite its soluble expression, the relatively low protein yield and the low *in vitro* solubility (see below) have hampered intensive engineering and application of *ta*CA. Although *ta*CA was expressed mostly in soluble forms regardless of the fusion tags, the highest expression level was attained when the NEXT tag was used **(Fig. 1d)**. Recently, *is*PETase has been extensively studied as a green biocatalyst that can degrade PET plastic under moderate temperature conditions (22-24). However, *is*PETase exhibits a low level of soluble expression in *E. coli* even at low temperature conditions, and it has not been demonstrated to have high-level soluble expression. The untagged *is*PETase was expressed almost exclusively in an insoluble form at 37 °C, as was the case for MBP and Fh8 fusion proteins. In contrast, surprisingly, NEXT-*is*PETase was expressed entirely in a soluble form **(Fig. 1e)**. Collectively, these results demonstrate that the NEXT tag is an exceptionally powerful enhancer not only for *in vivo* solubility but also for the production yield of passenger protein.

### Effect of the fusion tags on the *in vitro* solubility of purified proteins

To test the ability of solubility tags to promote the *in vitro* solubility of passenger proteins, *ta*CA and *is*PETase were further utilized for the test because these enzymes have low *in vitro* solubility and are susceptible to aggregation under low salt conditions (19, 24). The purified *ta*CA enzymes with different solubility tags were exposed to buffer solutions with or without salt supplementation, and any protein precipitates were separated from the supernatant by centrifugation and analyzed by SDS-PAGE. When the buffer was supplemented with 300 mM NaCl, only GST-*ta*CA showed a significant amount of precipitates, and the other proteins, including the untagged counterpart, remained soluble **(Fig. 2a)**. After the enzymes were placed in a low salt condition, GST-*ta*CA became completely insoluble, which was more severe than in the case of untagged *ta*CA **(Fig. 2b)**. Combined with the *in vivo* solubility results **(Fig. 1)**, this confirms that GST is not an effective tag for improving protein solubility (1). On the other hand, almost all *ta*CA enzymes were still in soluble forms when the other tags (MBP, Fh8, and NEXT) were used, demonstrating their effectiveness in improving the *in vitro* solubility of passenger proteins **(Fig. 2b)**. Interestingly, after undergoing changes in the composition and pH of buffer under low salt conditions, only MBP- or NEXT-tagged *ta*CA showed resistance to aggregation induced by changes in environmental conditions, indicating that both MBP and NEXT tags are superior to the Fh8 tag for enhancing *in vitro* solubility even under dynamic chemical environments **(Fig. 2c)**. Similar to *ta*CA, the poor *in vitro* solubility of *is*PETase under low salt conditions was successfully circumvented by fusion with the NEXT tag **(Fig. 2d)**.

**FIG 2.**
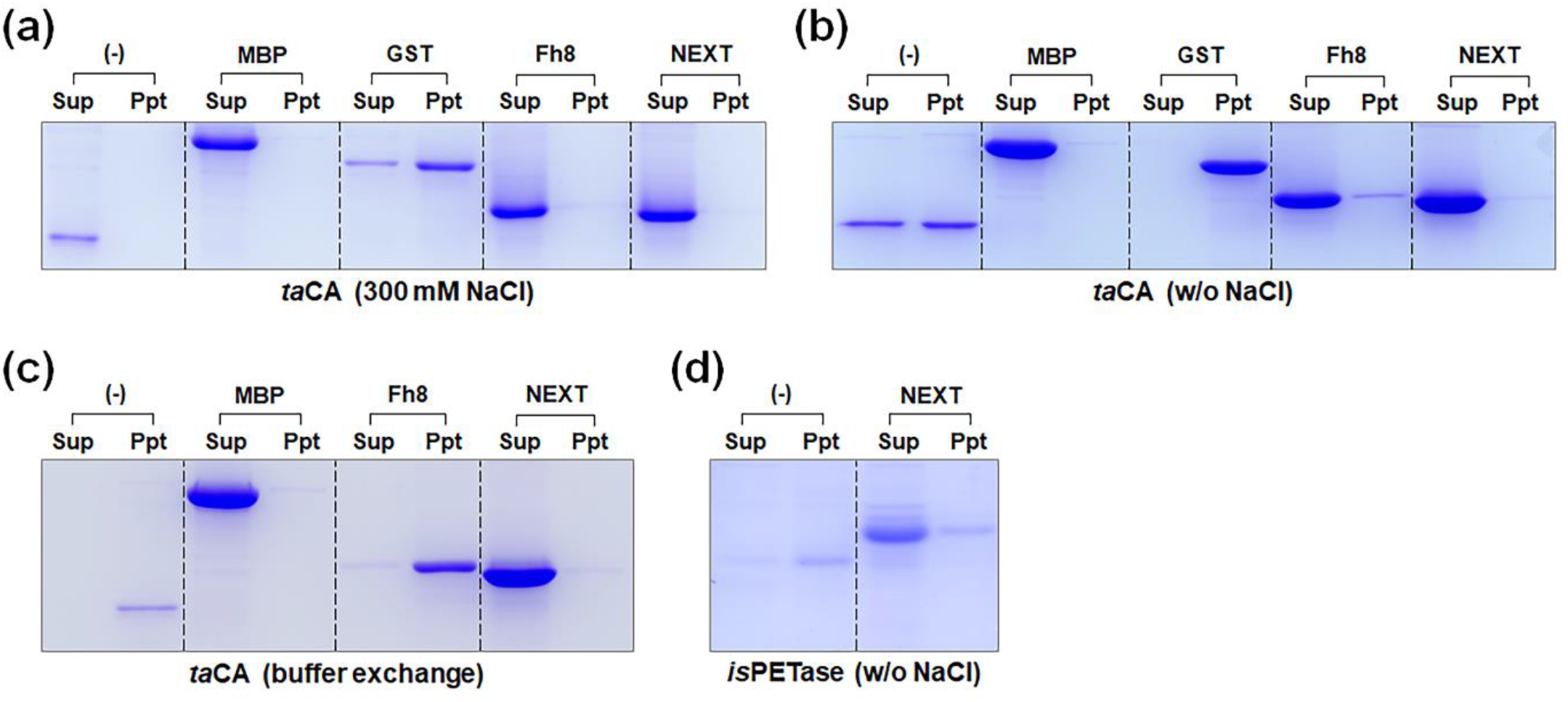
*In vitro* solubility of purified enzymes fused with different solubility tags. Coomassie blue-stained SDS-PAGE gel for the analysis of precipitation of (a) *ta*CA dialyzed against 20 mM phosphate buffer (pH 7.5) supplemented with 300 mM NaCl, (b) *ta*CA dialyzed against 20 mM phosphate buffer (pH 7.5), (c) *ta*CA after buffer exchange from 20 mM phosphate buffer (pH 7.5) to 20 mM Tris buffer (pH 8.3), and (d) *is*PETase dialyzed against 20 mM phosphate buffer (pH 7.5). Lane: Sup, supernatant after centrifugation; Ppt, precipitated protein pellet.

### Effect of the fusion tags on protein quality

Using the purified *ta*CA enzyme fused with the MBP, Fh8, or NEXT tag, the effect of the fusion tag on protein quality was investigated by examining the enzyme activity and stability, the two most important enzyme properties. The activity changes of *ta*CA caused by the fusion of Fh8 (27%) and NEXT (14%) were relatively marginal when compared with that of the MBP-*ta*CA, which showed an abnormally large increase in activity (115%) **(Fig. 3a)**. The bulky MBP tag might interfere with the function of *ta*CA more than the other small tags can. Similarly, the solubility tags affected the thermal stability of *ta*CA corresponding with their size, and no apparent decrease in stability was seen when the NEXT tag was used **(Fig. 3b)**. These results show that the smallest NEXT tag can be used as a noncleavable solubility tag that exerts only minimal influences on passenger proteins.

**FIG 3.**
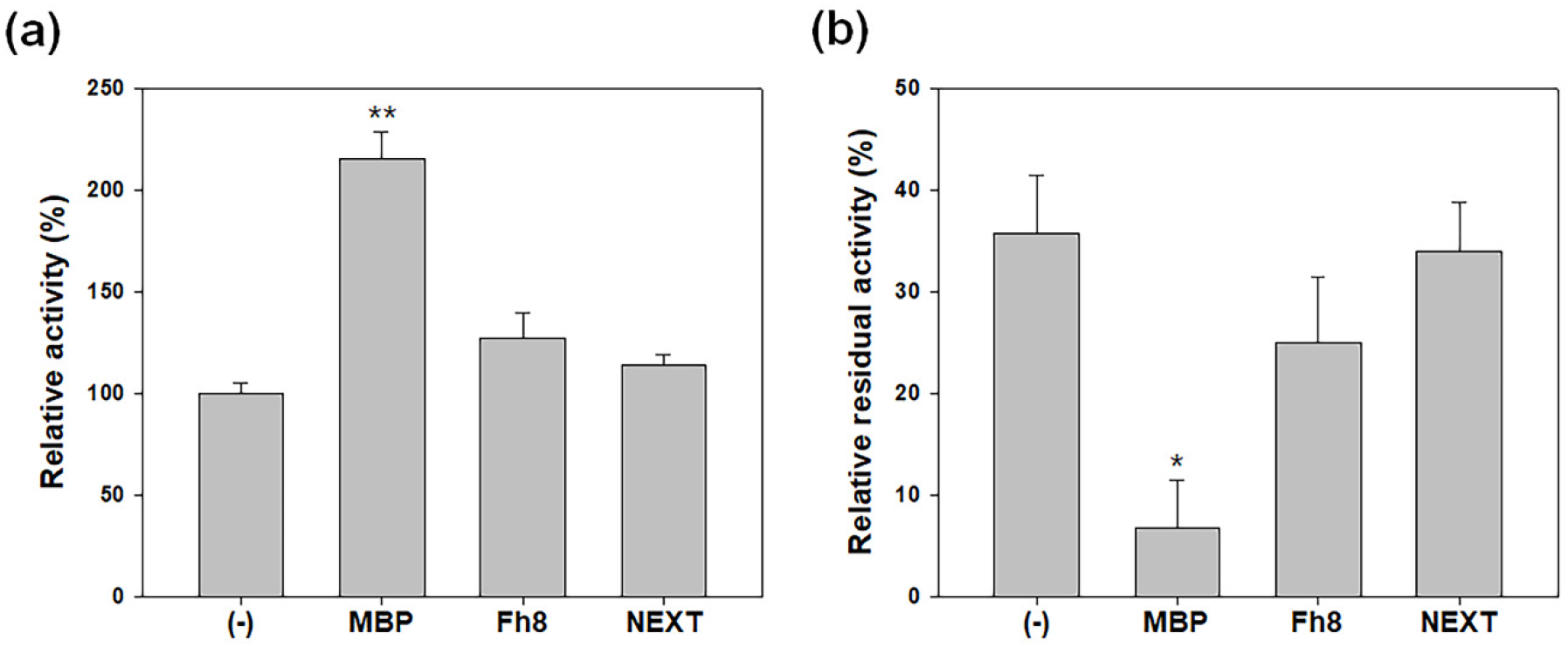
Activity and stability of purified *ta*CA fused with different tags. (a) Relative CO_2_ hydration activity of solubility tag-fused *ta*CA compared to the activity of *ta*CA without a tag. (b) Thermal stability of *ta*CA with or without a fusion tag. The enzyme activity was measured after incubation for 1 h at 90 °C, and the relative residual activity was obtained compared to the enzyme activity without heat treatment. Enzymes were prepared in 20 mM phosphate buffer (pH 7.5). Error bars represent standard deviations from two or three independent experiments. Asterisks indicate statistical significance of the tagged *ta*CA compared with the untagged enzyme determined by *t*-test (\**p* < .05, \*\**p* < .01).

From a different standpoint, biochemical studies of proteins can also benefit from the small size of the NEXT tag. Larger solubility tags are expected to have more sites for posttranslational modification, such as phosphorylation **(Table 1)**. For example, a candidate substrate for a kinase expressed with the fusion of MBP might be phosphorylated by the kinase within the MBP portion, not within the candidate substrate, leading to an incorrect conclusion that the candidate protein is actually a substrate for the kinase. In this situation, the NEXT tag with no potential phosphorylation site can be alternatively used.

### N-terminal truncation of the NEXT tag

To identify the part of the NEXT tag that is most responsible for solubility enhancement, the tag was roughly divided into three regions with similar sizes, and the sequential N-terminal truncation of the regions was studied **(Fig. 4a)**. First, *hm*CA, from which the NEXT tag originated, was expressed with the partial or full deletion of the N-terminal extension **(Fig. 4b)**. Full-length *hm*CA and the two partial deletion variants (ΔN19 and ΔN36) were expressed in soluble forms despite the different expression levels. However, when the N-terminal extension of *hm*CA was fully truncated, the protein was expressed in a form that was almost insoluble, as previously reported (9). This result clearly indicates that the C-terminal part of the NEXT tag (NEXT_C16_ peptide) is the part most responsible for the soluble expression of *hm*CA.

**FIG 4.**
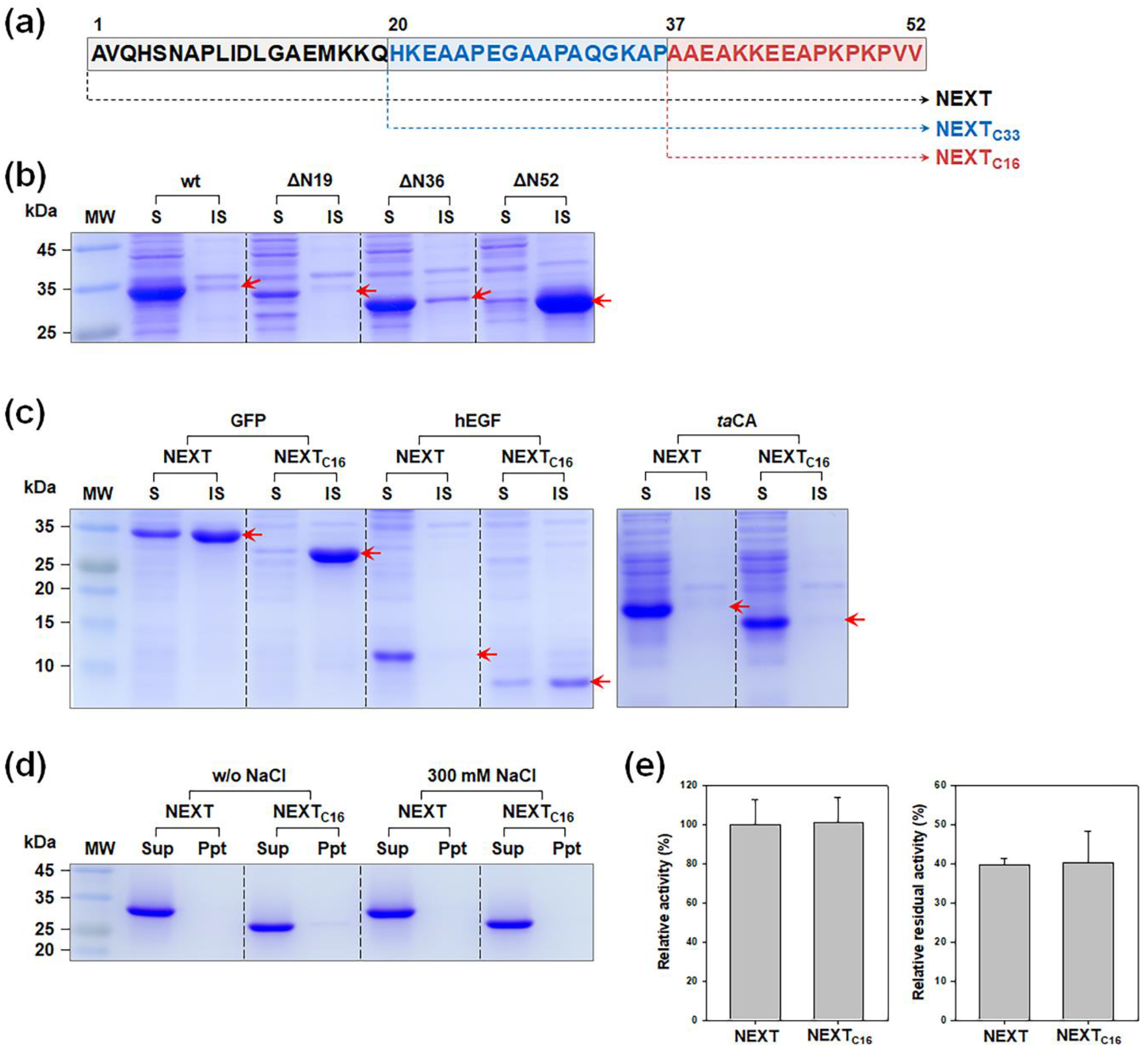
Effect of truncating the NEXT tag on the solubility of the target protein. (a) Design of truncation constructs. The sequence of the NEXT tag is presented along with the selected residue numbers. The first methionine was excluded for residue numbering. (b) *In vivo* solubility of N-terminal truncated *hm*CA variants. (c) *In vivo* solubility of the recombinant proteins (GFP, hEGF, and *ta*CA) fused with the NEXT or NEXT_C16_ tag. Proteins were expressed at 37 °C with 1 mM IPTG. (d) *In vitro* solubility of purified *ta*CA with the NEXT or NEXT_C16_ tag prepared in 20 mM phosphate buffer with or without 300 mM NaCl. Lane: MW, molecular mass marker; S, soluble fraction; IS, insoluble fraction; Sup, supernatant after centrifugation; Ppt, precipitated protein pellet. (e) Activity and stability of *ta*CA fused with the NEXT or NEXT_C16_ tag. *ta*CA enzymes were prepared in 20 mM phosphate buffer (pH 7.5). For the stability test, enzymes were incubated for 1 h at 90 °C. Error bars represent standard deviations from two independent experiments.

To test whether the 16 amino acid-length NEXT_C16_ can substitute the full-length NEXT tag, the expression patterns of hEGF, GFP, and *ta*CA fused to NEXT_C16_ were evaluated. Unfortunately, the soluble expressions of both hEGF and GFP were significantly hampered by the replacement of the NEXT tag with the NEXT_C16_ tag, suggesting that the N-terminal region of the NEXT tag is crucial for improving the *in vivo* solubility of the passenger protein **(Fig. 4c)**. In the case of *ta*CA, soluble expression of NEXT_C16_-*ta*CA was observed as expected, although the production yield appeared to be reduced compared to that of NEXT-*ta*CA **(Fig. 4c)**. Intriguingly, when the *in vitro* solubility of purified NEXT_C16_-*ta*CA was tested, no apparent protein precipitation was observed regardless of NaCl supplementation **(Fig. 4d)**. Additionally, the activity and stability of NEXT_C16_-*ta*CA were identical to those of NEXT-*ta*CA **(Fig. 4e)**. These results show that in contrast to the *in vivo* solubility result, the high *in vitro* solubility of passenger proteins can be retained by using the NEXT_C16_ region alone instead of the full-length NEXT tag. The use of a very short NEXT_C16_ tag might be beneficial, *e*.*g*., for the immobilization of the target enzyme onto a solid matrix with a limited surface area to maximize the immobilization yield and overall catalytic efficiency of biocatalysts (25).

### Intrinsic disorder of the NEXT tag

The mechanisms of solubility enhancement by various solubility tags are still not clear, and there seems to be multiple routes for the promoted solubility (4). The *in vivo* solubility results **(Fig. 1)** did not fit into the solubility prediction by the modified Wilkinson−Harrison model **(Table S1)** (26). Although machine learning-based SOLpro (27) predicted the solubility patterns of fusion proteins more accurately, it could still not discriminate the NEXT tag from the others, especially for *is*PETase fusions **(Table S1, Fig. 1e)**. Protein acidity is known to be one of the determinants of solubility (28, 29), which cannot explain the remarkable enhancement of solubility by the NEXT tag, which possesses a net positive charge **(Table 1)**. The classical structure-function paradigm of proteins has been challenged by the concept of intrinsically disordered proteins (IDPs). IDPs exist as highly dynamic structural ensembles with undefined three-dimensional structures (30). It has been proposed that an IDP region within a protein can act as an intramolecular entropic bristle (EB) (31, 32). The EB domain is expected to have an extended conformation, and by thermally driven random motion, it can occupy a significantly large space around the protein molecule (33). By entropically excluding neighboring molecules, EB can prevent protein aggregation, thus leading to improved protein solubility.

Sequence-based prediction showed that the NEXT tag is highly disordered, whereas the MBP, GST, and Fh8 tags possess low disorder propensities **(Fig. 5a)**. Its low hydrophobicity, a measure that is correlated with protein disorder (34), also distinguishes the NEXT tag from the other tags **(Table 1)**. The C-terminal region of the NEXT tag was predicted to be the most disordered **(Fig. 5a)**, which corresponds to the NEXT_C16_ region that was crucial for improving solubility **(Fig. 4)**. The NEXT tag was separately expressed and purified **(Fig. 5b)**. The circular dichroism (CD) spectrum of the purified NEXT tag coincided with that of a random coil without any secondary structural element **(Fig. 5c)** (35). These results strongly suggest that the NEXT tag is an IDP that can improve the solubility of passenger proteins as a function of EB.

**FIG 5.**
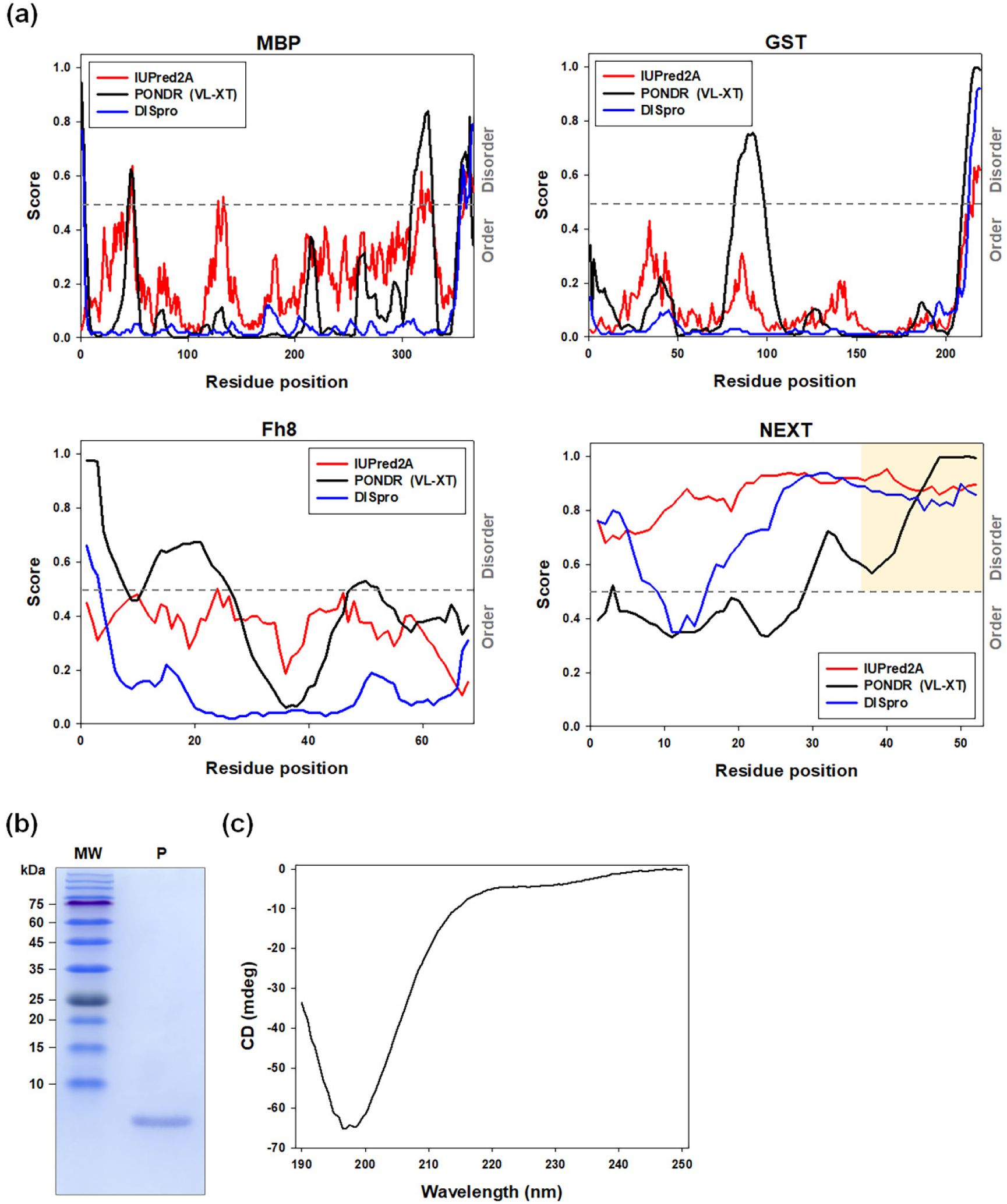
Intrinsic disorder propensities of solubility tags. (a) Position-dependent prediction of disordered regions by three different methods (IUPred2A, PONDR, and DISpro). The region corresponding to NEXT_C16_ is highlighted in a yellow box. (b) Purified NEXT tag with a C-terminal His_6_-tag (6.5 kDa) analyzed by SDS-PAGE. Lane: MW, molecular mass marker; P, purified protein. (c) Circular dichroism (CD) spectrum of purified NEXT tag in the far-UV region.

As previously demonstrated, the fusion of the NEXT tag can prevent protein aggregation both *in vitro* and *in vivo* **(Fig. 6)**. A fully folded protein with low *in vitro* solubility is prone to aggregation **(Fig. 6a)**, which can be circumvented by the fusion of EB **(Fig. 6b)**. The accumulation of overexpressed, fully folded protein with low *in vitro* solubility can result in protein aggregation in the cytosol **(Fig. 6c)**. This apparently low *in vivo* solubility can also be overcome by the utilization of EB **(Fig. 6d)**. The accumulation and subsequent aggregation of partially folded proteins before the completion of folding is another cause of low *in vivo* solubility **(Fig. 6e)**. The interaction between the folding intermediates can be reduced with N-terminal fusion of EB, facilitating correct protein folding **(Fig. 6f)**.

**FIG 6.**
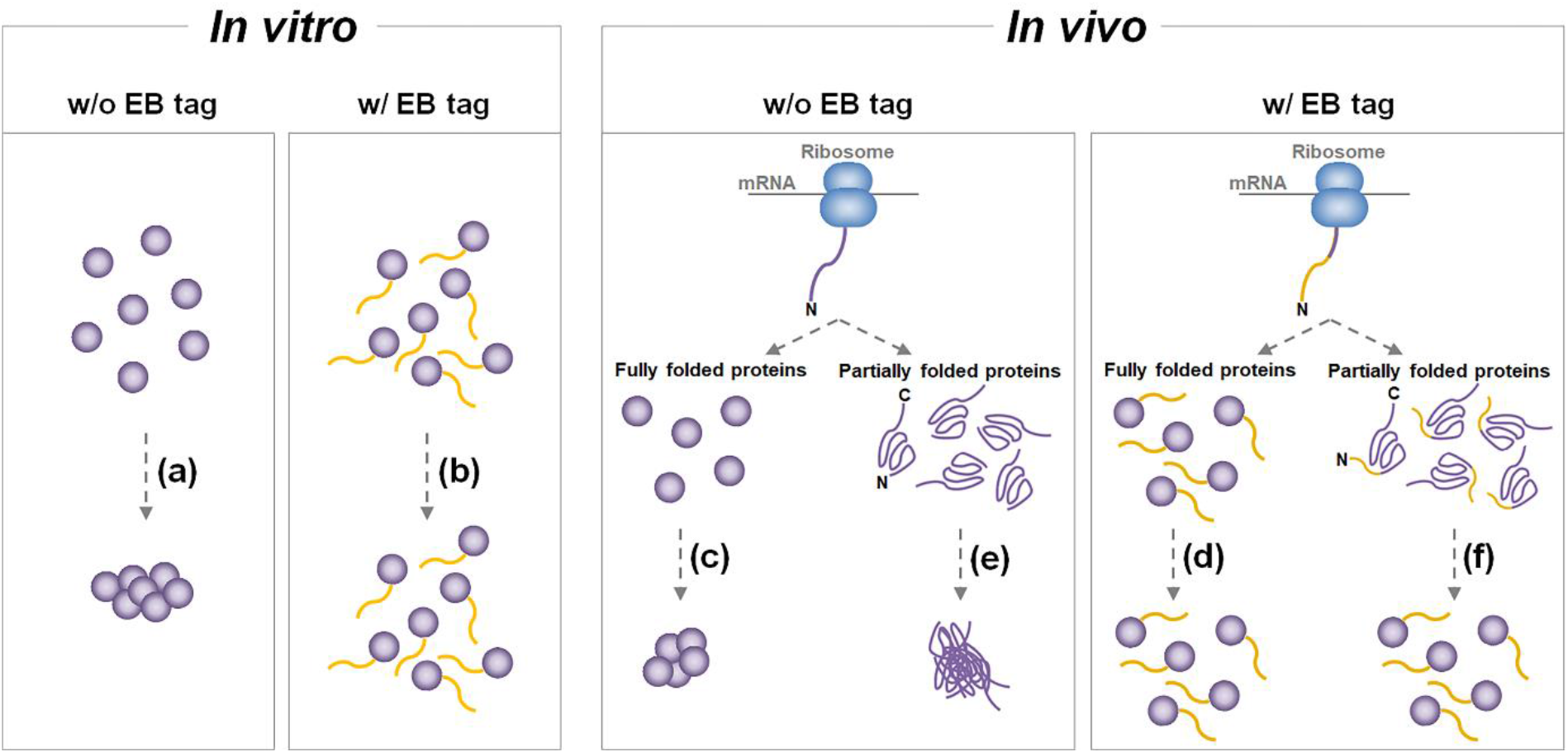
Solubility enhancement by the fusion of entropic bristle (EB) (a, b) A protein with low *in vitro* solubility aggregates in its folded state and after the fusion of EB, the protein can remain soluble. (c, d) The accumulation of fully folded protein with low *in vitro* solubility can lead to protein aggregation in the cytosol, leading to protein expression that seems to be insoluble. This situation can also be circumvented by the fusion of EB. (e, f) Low *in vivo* solubility during recombinant protein overexpression can occur by the aggregation of partially folded proteins, which can be overcome by the N-terminal fusion of EB, preventing the interaction between the folding intermediates and thus allowing protein folding to complete.

In conclusion, the successful use of a small-sized NEXT tag was demonstrated to improve both the *in vivo* and *in vitro* solubilities of the selected recombinant proteins. Because the degree of solubility enhancement by EB fusion appeared to depend on the length of the tag (31), a more powerful IDP-based solubility tag should be artificially engineered by optimizing not only the amino acid sequence but also the length of the tag. Further experimental analyses using a variety of potential EB proteins, including the NEXT tag, will provide insight into the engineering principles for the *de novo* design of IDP-based solubility tags customized for each passenger protein.

## MATERIALS AND METHODS

### Construction of expression vectors

The strains, plasmids, and primers used in this study are listed in **Table S2**. The *E. coli* TOP10 strain (Thermo Fisher Scientific, USA) was used for gene cloning. *E. coli* was routinely cultured in Luria-Bertani (LB) medium supplemented with appropriate antibiotics (10 μg/mL streptomycin or 50 μg/mL ampicillin) at 37 °C in a shaking incubator (Jeiotech, Korea). The genes for MBP, GST, NEXT, GFP and *ta*CA were cloned by polymerase chain reaction (PCR) using pMAL-c5X (New England Biolabs, USA), pGEX-4T-1 (GE Healthcare, USA), pET-hmCA (9), pTH-GFP (36), and pET-taCA (19) as the templates. The primers for the solubility tags contain the sequence for flexible linker (GGGGS)2 along with *Nde*I and *Nco*I restriction sites. The PCR fragments were cloned into the pGEM-T Easy vector (Promega, USA) and the amplified sequences were confirmed by direct sequencing. The genes for the Fh8 tag (GenBank accession number: AF213970), hEGF (GenBank accession number: M15672) and *is*PETase (GenBank accession number: 6EQD_A) were chemically synthesized along with the linker sequence (only for Fh8) and the restriction sites (Genotech, Korea). The genes were subcloned into pET-22b(+) (Novagen, USA). All of the recombinant genes have a hexahistidine (His_6_)-tag sequence at their 3’ termini provided by the parent vector.

### Expression of recombinant proteins

Recombinant *E. coli* BL21(DE3) strains transformed with the constructed vectors were incubated in LB medium with 50 μg/mL ampicillin at 37 °C in the shaking incubator. Protein expression was induced by adding isopropyl-β-D-thiogalactopyranoside (IPTG; Duchefa Biochemie, Netherlands) to a final concentration of 1 mM (at 37 °C) or 10 μM (at 25 °C) when the optical density at 600 nm (OD_600_) reached 0.6-0.8. For the expression of *ta*CA variants, 0.1 mM ZnSO_4_ (Junsei, Japan) was also added to the culture medium. After cultivation for 10 h at 37 °C or for 20 h at 25 °C, the cells were collected by centrifugation at 4 °C and 4,000 g for 10 min. The cells were resuspended in lysis buffer (50 mM sodium phosphate, 300 mM NaCl, and 10 mM imidazole; pH 8.0) and disrupted by an ultrasonic dismembrator (Sonics and Materials, USA) in ice water. After centrifugation of the lysate at 4 °C and 10,000*g* for 10 min, the supernatant was removed, and the soluble fraction (S) was designated; the remaining debris was designated the insoluble fraction (IS).

### Purification of recombinant proteins

The soluble fraction of the cell lysate was mixed with Ni^2+^-nitrilotriacetic acid agarose beads (Qiagen, USA), and the His_6_-tagged recombinant proteins were purified by immobilized metal affinity chromatography (IMAC) according to the manufacturer’s instructions. The enzymes were eluted using elution buffer (50 mM sodium phosphate, 300 mM NaCl, and 250 mM imidazole; pH 8.0). The eluates were thoroughly dialyzed against 20 mM sodium phosphate buffer (pH 7.5) with or without 300 mM NaCl. After dialysis was completed, the protein precipitates (Ppt) were removed by centrifugation at 4 °C and 10,000 g for 10 min. The supernatants (Sup) were used for subsequent activity and stability tests. In some cases, the enzyme buffer was further exchanged with 20 mM Tris-sulfate buffer (pH 8.3).

### Protein analyses

For protein quantification, the purified enzyme was denatured in denaturing buffer (6 M guanidine hydrochloride GuHCl/20 mM sodium phosphate buffer, pH 7.5), and the absorbance of the denatured protein was measured at 280 nm. The protein concentration was determined using the measured absorbance and the molar extinction coefficient at 280 nm for each protein calculated by ProtParam (http://web.expasy.org/protparam/) (37). Proteins were separated and visualized by sodium dodecyl sulfate-polyacrylamide gel electrophoresis (SDS-PAGE) followed by Coomassie Brilliant Blue R-250 (Bio-Rad, USA) staining. The percentage of soluble expression was estimated by densitometric analysis of the band intensities of soluble and insoluble fractions on the protein gel performed using ImageJ.

### Activity and stability test for *ta*CA variants

CA activity was measured by a colorimetric CO_2_ hydration assay (38, 39). The purified enzyme in 20 mM phosphate buffer (pH 7.5) was diluted to a concentration of 1 μM. Ten microliters of sample was added to a disposable cuvette containing 600 μL of 20 mM Tris buffer (pH 8.3) supplemented with 100 μM phenol red. The reaction was performed at 4 °C inside the spectrometer by adding 400 μL of CO_2_-saturated deionized water prepared in ice-cold water. The absorbance change was monitored at 570 nm. The time (*t*) required for the absorbance to drop from 1.2 (corresponding to pH 7.5) to 0.18 (corresponding to pH 6.5) was obtained. The time (*t*_0_) for the uncatalyzed reaction was also measured by adding a corresponding blank buffer instead of an enzyme sample. The Wilbur-Anderson unit was calculated as (*t*_0_− *t*)/*t*. For the stability test, the enzyme sample was incubated for 1 h at 90 °C and the residual enzyme activity was measured. Relative residual activity was calculated based on the activity of the untreated sample.

### Circular dichroism spectroscopy

Circular dichroism (CD) spectrum was recorded on a CD spectropolarimeter (Jasco, Japan). The purified solution of the NEXT tag in 20 mM phosphate buffer (pH 7.5) was scanned in a quartz crystal cuvette with a 2 mm path length (Hellma Analytics, Germany) for the far-UV region (190-250 nm) at 25 °C. Based on the CD spectrum, secondary structural elements were analyzed using BeStSel (40).

### *In silico* calculations

Protein parameters including amino acid length, molecular weight, net charge and pI, were calculated by ProtParam (37). Phosphorylation sites were predicted by NetPhos 3.1 (http://www.cbs.dtu.dk/services/NetPhos/) (41). Kyte-Doolittle hydropathicity was calculated by ProtScale (https://web.expasy.org/protscale/) using a window size of 5, and the values were averaged to obtain a mean hydropathy (37, 42). Sequence-based prediction of protein solubility was performed by the modified Wilkinson−Harrison method (26) and SOLpro (http://scratch.proteomics.ics.uci.edu/) (27). Disordered protein regions were predicted by IUPred2A (https://iupred2a.elte.hu/) (43), PONDR (http://www.pondr.com/) (44) and DISpro (http://scratch.proteomics.ics.uci.edu/) (45).

## ACKNOWLEDGMENTS

This work was supported by the Korea Institute of Energy Technology Evaluation and Planning (KETEP) grant (20182010600430) funded by the Ministry of Trade, Industry, and Energy, Korea, and by the National Research Foundation grants (NRF-2020M3A9I5037642, NRF-2021R1F1A1057310 and NRF-2021R1A5A8029490) funded by the Ministry of Science and ICT, Korea.

## REFERENCES

1. Rosano GL, Ceccarelli EA. 2014. Recombinant protein expression in Escherichia coli: advances and challenges. Front Microbiol 5:172.

2. Sorensen HP, Mortensen KK. 2005. Soluble expression of recombinant proteins in the cytoplasm of Escherichia coli. Microb Cell Fact 4:1.

3. Singh A, Upadhyay V, Upadhyay AK, Singh SM, Panda AK. 2015. Protein recovery from inclusion bodies of Escherichia coli using mild solubilization process. Microb Cell Fact 14:41.

4. Costa S, Almeida A, Castro A, Domingues L. 2014. Fusion tags for protein solubility, purification, and immunogenicity in Escherichia coli: the novel Fh8 system. Front Microbiol 5:63.

5. Esposito D, Chatterjee DK. 2006. Enhancement of soluble protein expression through the use of fusion tags. Curr Opin Biotechnol 17:353–358.

6. Yang J, Han YH, Im J, Seo SW. 2021. Synthetic protein quality control to enhance full-length translation in bacteria. Nat Chem Biol 17:421–427.

7. Zhou P, Wagner G. 2010. Overcoming the solubility limit with solubility-enhancement tags: successful applications in biomolecular NMR studies. J Biomol NMR 46:23–31.

8. Seijsing J, Lindborg M, Hoiden-Guthenberg I, Bonisch H, Guneriusson E, Frejd FY, Abrahmsen L, Ekblad C, Lofblom J, Uhlen M, Graslund T. 2014. An engineered affibody molecule with pH-dependent binding to FcRn mediates extended circulatory half-life of a fusion protein. Proc Natl Acad Sci USA 111:17110–17115.

9. Jo BH, Im SK, Cha HJ. 2018. Halotolerant carbonic anhydrase with unusual N-terminal extension from marine Hydrogenovibrio marinus as novel biocatalyst for carbon sequestration under high-salt environments. J CO2 Util 26:415–424.

10. Trevino SR, Scholtz JM, Pace CN. 2008. Measuring and increasing protein solubility. J Pharm Sci 97:4155–4166.

11. Golovanov AP, Hautbergue GM, Wilson SA, Lian LY. 2004. A simple method for improving protein solubility and long-term stability. J Am Chem Soc 126:8933–8939.

12. Kapust RB, Waugh DS. 1999. Escherichia coli maltose-binding protein is uncommonly effective at promoting the solubility of polypeptides to which it is fused. Protein Sci 8:1668–1674.

13. Harper S, Speicher DW. 2008. Expression and purification of GST fusion proteins. Curr Protoc Protein Sci 52:6.6.1-6.6.28.

14. Choi JS, Leong KW, Yoo HS. 2008. In vivo wound healing of diabetic ulcers using electrospun nanofibers immobilized with human epidermal growth factor (EGF). Biomaterials 29:587–596.

15. Kang YS, Song JA, Han KY, Lee J. 2015. Escherichia coli EDA is a novel fusion expression partner to improve solubility of aggregation-prone heterologous proteins. J Biotechnol 194:39–47.

16. Zheng XM, Wu X, Fu XL, Dai DP, Wang FH. 2016. Expression and purification of human epidermal growth factor (hEGF) fused with GB1. Biotechnol Biotechnol Equip 30:813–818.

17. Kim YS, Lee HJ, Han MH, Yoon NK, Kim YC, Ahn J. 2021. Effective production of human growth factors in Escherichia coli by fusing with small protein 6HFh8. Microb Cell Fact 20:9.

18. Tsien RY. 1998. The green fluorescent protein. Annu Rev Biochem 67:509–544.

19. Jo BH, Seo JH, Cha HJ. 2014. Bacterial extremo-α-carbonic anhydrases from deep-sea hydrothermal vents as potential biocatalysts for CO_2_ sequestration. J Mol Catal B-Enzym 109:31–39.

20. Parra-Cruz R, Lau PL, Loh HS, Pordea A. 2020. Engineering of Thermovibrio ammonificans carbonic anhydrase mutants with increased thermostability. J CO2 Util 37:1–8.

21. Nguyen TKM, Ki MR, Son RG, Pack SP. 2019. The NT11, a novel fusion tag for enhancing protein expression in Escherichia coli. Appl Microbiol Biotechnol 103:2205–2216.

22. Yoshida S, Hiraga K, Takehana T, Taniguchi I, Yamaji H, Maeda Y, Toyohara K, Miyamoto K, Kimura Y, Oda K. 2016. A bacterium that degrades and assimilates poly(ethylene terephthalate). Science 351:1196–1199.

23. Son HF, Cho IJ, Joo S, Seo H, Sagong HY, Choi SY, Lee SY, Kim KJ. 2019. Rational protein engineering of thermo-stable PETase from Ideonella sakaiensis for highly efficient PET degradation. ACS Catal 9:3519–3526.

24. Taniguchi I, Yoshida S, Hiraga K, Miyamoto K, Kimura Y, Oda K. 2019. Biodegradation of PET: current status and application aspects. ACS Catal 9:4089–4105.

25. Sheldon RA, van Pelt S. 2013. Enzyme immobilisation in biocatalysis: why, what and how. Chem Soc Rev 42:6223–6235.

26. Davis GD, Elisee C, Newham DM, Harrison RG. 1999. New fusion protein systems designed to give soluble expression in Escherichia coli. Biotechnol Bioeng 65:382–388.

27. Magnan CN, Randall A, Baldi P. 2009. SOLpro: accurate sequence-based prediction of protein solubility. Bioinformatics 25:2200–2207.

28. Su Y, Zou Z, Feng S, Zhou P, Cao L. 2007. The acidity of protein fusion partners predominantly determines the efficacy to improve the solubility of the target proteins expressed in Escherichia coli. J Biotechnol 129:373–382.

29. Kramer RM, Shende VR, Motl N, Pace CN, Scholtz JM. 2012. Toward a molecular understanding of protein solubility: increased negative surface charge correlates with increased solubility. Biophys J 102:1907–1915.

30. Uversky VN. 2019. Intrinsically disordered proteins and their “mysterious” (meta)physics. Front Phys 7:10.

31. Santner AA, Croy CH, Vasanwala FH, Uversky VN, Van YY, Dunker AK. 2012. Sweeping away protein aggregation with entropic bristles: intrinsically disordered protein fusions enhance soluble expression. Biochemistry 51:7250–7262.

32. Grana-Montes R, Marinelli P, Reverter D, Ventura S. 2014. N-terminal protein tails act as aggregation protective entropic bristles: the SUMO case. Biomacromolecules 15:1194–1203.

33. Hoh JH. 1998. Functional protein domains from the thermally driven motion of polypeptide chains: a proposal. Proteins 32:223–228.

34. Uversky VN, Gillespie JR, Fink AL. 2000. Why are “natively unfolded” proteins unstructured under physiologic conditions? Proteins 41:415–427.

35. Srinivasan N, Bhagawati M, Ananthanarayanan B, Kumar S. 2014. Stimuli-sensitive intrinsically disordered protein brushes. Nat Commun 5:5145.

36. Cha HJ, Wu CF, Valdes JJ, Rao G, Bentley WE. 2000. Observations of green fluorescent protein as a fusion partner in genetically engineered Escherichia coli: monitoring protein expression and solubility. Biotechnol Bioeng 67:565–574.

37. Wilkins MR, Gasteiger E, Bairoch A, Sanchez JC, Williams KL, Appel RD, Hochstrasser DF. 1999. Protein identification and analysis tools in the ExPASy server. Methods Mol Biol 112:531–552.

38. Jo BH, Moon H, Cha HJ. 2020. Engineering the genetic components of a whole-cell catalyst for improved enzymatic CO_2_ capture and utilization. Biotechnol Bioeng 117:39–48.

39. Wilbur KM, Anderson NG. 1948. Electrometric and colorimetric determination of carbonic anhydrase. J Biol Chem 176:147–154.

40. Micsonai A, Wien F, Bulyaki E, Kun J, Moussong E, Lee YH, Goto Y, Refregiers M, Kardos J. 2018. BeStSel: a web server for accurate protein secondary structure prediction and fold recognition from the circular dichroism spectra. Nucleic Acids Res 46:W315–W322.

41. Blom N, Gammeltoft S, Brunak S. 1999. Sequence and structure-based prediction of eukaryotic protein phosphorylation sites. J Mol Biol 294:1351–1362.

42. Kyte J, Doolittle RF. 1982. A simple method for displaying the hydropathic character of a protein. J Mol Biol 157:105–132.

43. Meszaros B, Erdos G, Dosztanyi Z. 2018. IUPred2A: context-dependent prediction of protein disorder as a function of redox state and protein binding. Nucleic Acids Res 46:W329–W337.

44. Romero P, Obradovic Z, Li X, Garner EC, Brown CJ, Dunker AK. 2001. Sequence complexity of disordered protein. Proteins 42:38–48.

45. Cheng J, Sweredoski MJ, Baldi P. 2005. Accurate prediction of protein disordered regions by mining protein structure data. Data Min Knowl Disc 11:213–222.

